# Neuropilin-1 in CGRP Afferents Is Required for VEGFA-Driven Pain and Baseline Nociceptive Sensitivity

**DOI:** 10.64898/2026.05.06.723195

**Authors:** Shilang Xiao, Heather N. Allen, Olivia L. Babyok, Santiago Loya Lopez, Stephanie A. Fulton, Tyler S. Nelson, Rajesh Khanna, Jami L. Saloman

**Affiliations:** Department of Gastroenterology, The Third Xiangya Hospital, Central South University, School of Medicine, Changsha, Hunan 410013, China. Xiangya Scholars Program, University of Pittsburgh, Pittsburgh, PA, USA; Department of Medicine, University of Pittsburgh, Pittsburgh, PA, USA; Department of Pharmacology and Therapeutics, McKnight Brain Institute, and Center for Advanced Pain Therapeutics and Research (CAPToR), College of Medicine, University of Florida, Gainesville, Florida, USA; Pittsburgh Center for Pain Research, University of Pittsburgh, Pittsburgh, PA, USA; Department of Neurobiology and Center for Neuroscience, University of Pittsburgh, Pittsburgh, PA, USA; Physician Scientist Training Program (PSTP), University of Pittsburgh, Pittsburgh, PA, USA

**Keywords:** Neuropilin-1, VEGFA, peptidergic neuron

## Abstract

Neuropilin-1 (NRP1) is a single pass transmembrane glycoprotein that can form a receptor complex with several tyrosine kinase receptors, including the vascular endothelial growth factor (VEGF) receptor. Previous studies have reported that binding of VEGFA to this receptor complex elicits mechanical allodynia and thermal hyperalgesia through potentiation of voltage-gated sodium and calcium channel activity. We find that *Nrp1* mRNA and protein is widely distributed in naïve mouse and rat DRG neurons, including peptidergic afferents. A CGRPcreER: NRP1^fl/fl^ transgenic mice were generated to investigate the role of peptidergic NRP1 in basal nociception. Following *in vivo* loss of NRP1, mice are hyposensitive to both noxious heat and mechanical stimuli. Under normal conditions, VEGFA elicits mechanical hypersensitivity, an effect that was absent in our NRP1 knockout mouse. Furthermore, VEGFA induced neuronal hyperexcitability was lost in CGRP expressing neurons isolated from this NRP1 knockout mouse. This study validates the NRP1 knockout mouse and confirms previous findings that VEGFA, often released during pathological pain conditions, requires peptidergic NRP1. Interestingly, we find that in the absence of ongoing injury or inflammation, peptidergic NRP1 regulates basal nociception and pain-like behaviors.

**Perspective:** NRP1 is expressed in sensory neurons including the peptidergic subpopulation. Genetic deletion of NRP1 in healthy adults alters nociception without altering innervation; NRP1 knockout mice are hyposensitive to noxious heat and mechanical stimuli, but lose sensitivity to VEGFA, confirming it is a therapeutic target for growth factor mediated pain conditions.

## Introduction

Neuropilin-1 (NRP1) is a single pass transmembrane glycoprotein expressed in neurons, vascular endothelial cells, and multiple types of immune cells, including regulatory T cells (Treg), CD8 T cells, and dendritic cells.[14; 18; 24; 36; 37] NRP1 acts as a co-receptor and forms complexes with other tyrosine kinase receptors (TRKs), including plexin A1, plexin A2, VEGFR1, and VEGFR2. These receptor complexes bind the class III Semaphorin family members (Sema3, ligands of plexins) or vascular endothelial growth factors (VEGFs, ligands of VEGFRs) to transduce biological responses related to neuronal development, axonal guidance, angiogenesis, and vascularization.[28]

NRP1 has been described as an axon-guidance protein receptor due to its ability to mediate axonal growth in neuronal development via cooperation with other axon guidance cues, including SEMA3A and VEGF165. [31; 41] However, recent studies have identified a novel role for NRP1 in the context of pain. Activation of NRP1 signaling evoked by VEGFA increases voltage-gated sodium and calcium channel activity in dorsal root ganglia (DRG), leading to mechanical allodynia and thermal hyperalgesia in rats. [22; 33] In concurrence with this pronociceptive role for NRP1, downregulation of NRP1 in DRG effectively blocks spared nerve ligation-induced neuropathic pain in rats.[12] Given the involvement of VEGFA in pathological conditions such as osteoarthritis [34], endometriosis [23], and cancer [9], and the newly defined role of NRP1 in pain modulation, the VEGFA/NRP1 signaling cascade may be a target for effective pain control in different diseases.

Previous studies as well as our studies have demonstrated that NRP1 is widely distributed in sensory DRG neurons. However, previous studies applied CRISPR/Cas9 editing to delete NRP1 in all DRG neurons to investigate the effect of NRP1 on ion channel activity and pain behavior[12]; thus, the specific subset of DRG neuron responsible for NRP1-mediated pain hyposensitivity remains unknown. Most sensory neurons innervating certain targets (e.g. colon, pancreas) express some amount of *calca* mRNA (gene encoding CGRP), regardless of cluster assignment.[21; 35] CGRP-expressing sensory neurons, a major peptidergic subpopulation, are key regulators of neurogenic inflammation and nerve injury related pain. For instance, CGRP afferents mediate chronic pancreatitis pain in a nerve growth factor (NGF) dependent manner [15]; interestingly, NGF and TGFβ are also ligands for NRP1:TRK complexes.[26] These growth factors are highly upregulated in multiple cancers, painful diseases in which CGRP-expressing sensory neurons have also been directly implicated.[20; 26] Therefore, we established a new transgenic mouse model (CGRPcreER:NRP1^fl/fl^) to specifically interrogate the role of NRP1 in this peptidergic population, creating a tool for future studies on painful conditions such as cancer and pancreatitis. To validate and characterize this new transgenic model, we investigated the effect of NRP1 deletion in CGRP-sensory neurons on basal nociceptive behaviors, VEGFA-mediated allodynia and excitability. In this study, we demonstrate that NRP1 expression is enriched in peptidergic neurons, and that targeted deletion of NRP1 within these neurons significantly attenuates VEGFA-induced neuronal excitability, markedly reduces mechanical and thermal sensitivity, and effectively abolishes VEGFA-mediated allodynia.

## Materials and Methods

### Mice

Male and female (7-30 weeks old) C57BL/6 laboratory mice were housed under AAALAC-accredited conditions at the University of Pittsburgh or University of Florida in accordance with guidelines of the Institutional Animal Care and Use Committees and the National Institutes of Health Guide for the Care and Use of Laboratory Animals. In this study, CGRPcreER:NRP1^fl/fl^ mice were developed by crossing CGRPcreER mice (gift from Dr. Pao-Tien Chuang, UCSF) with NRP1^fl/fl^ (gift from Dr. Dario Vignali, University of Pittsburgh) to induce specific deletion of NRP1 in CGRP positive neurons. The TRPV1cre strain (Jackson Laboratory, Stock #017769) was crossed to the NRP1^fl/fl^ to generate peptidergic sensory neuron specific NRP1 deletion. In a subset of experiments, mice were further crossed with the Ai9 Cre-dependent reporter line (Jackson Laboratory stock #007909) to drive tdTomato expression, enabling reliable identification of knockout neurons in vitro.

### Rats

Adult male and female Sprague-Dawley rats (100-200g) were purchased from Charles River Laboratories and housed in New York University’s Kriser Dental Facility on a 12-hour light cycle with *ad libitum* access to food and water. All procedures were accordance with New York University’s Institutional Care and Use Committee and National Institutes of Health Guide for the Care and Use of Laboratory Animals.

### Tamoxifen

Mice received intraperitoneal injection of tamoxifen (Sigma-Aldrich, Cat: T5648-5G, 100mg /kg in corn oil) for 5 consecutive days for conditional deletion of NRP1. Mice were used for subsequent experiments at least 14 days after the final tamoxifen injection.

### Basescope with Immunofluorescence

Mice were euthanized with overdose isoflurane and perfused transcardially with 4% PFA. The L4DRGs were dissected and post-fixed in 4% PFA for two hours. The DRGs were cryoprotected in 25% sucrose buffer at 4°C, embedded in OCT, and frozen in -80°C until cryosectioning. The tissues were serially sectioned to 14 µm thickness and mounted on Superfrost Plus slides. Slides were bathed in distilled water to remove OCT and incubated with hydrogen peroxide for 2 min. Slides were washed again and air dried at room temperature overnight. Slides were treated with protease III for 20 min at 40°C within HybEZ Oven prior to incubation of C1 probe targeting NRP1 exon2 for 2 hours. Signal amplification and detection reagents (Cat. #323910, ACDBio) were applied sequentially and incubated in AMP1, AMP2, AMP3, AMP4, AMP5, AMP6, AMP7, AMP8 and RED solution reagents, for 30min, 30min, 15min, 30min, 30min, 15min, 30min, 15min, 10min, respectively. The slides were then washed with PBST buffer and blocked in 4% normal donkey serum, 0.25% triton, and 1X Phosphate Buffered Saline for 1 hour. Slides were incubated in primary antibody against CGRP, TH, NF200, or PGP9.5 diluted in PBST buffer overnight, rinsed three times in PBST buffer for 5 minutes each, and incubated in secondary antibody for 1 hour (Anti-Rabbit-Cy3 or Anti-rabbit-Cy5 1:500, Jackson ImmunoResearch). Finally, the slides were rinsed with PBST buffer three times and coverslipped with prolong golden antifade medium (Cat# P36941, Thermo Scientific). Detailed antibody information including RRIDs is provided in **supplementary table 1**. Cells with at least 15 puncta were considered *Nrp1* positive. Antibody staining was determined as positive using the threshold tool in Image J combined with controls. Controls included positive and negative control slides provided by ACDBio as well as performing protocols in the absence of the probe, primary, and/or secondary antibody. Three to eight sections per individual mouse were analyzed to calculate the percentage of different neuron subsets. The final images were produced in Adobe Illustrator 2025.

### RNAscope Multiplex v2 Fluorescent Assay

Rats were deeply anesthetized with isoflurane and euthanized through decapitation. Lumbar (L4 and L5) dorsal root ganglia (DRG) were rapidly extracted through blunt dissection and frozen in optimal cutting temperature (OCT) on dry ice. Tissues were stored at -80°C until cryosectioning. DRG tissue was sectioned at 18 μm, directly mounted on Superfrost Plus Microscope slides, and air dried overnight at room temperature (RT). The next morning slides underwent pretreatment for RNAscope fluorescence in situ hybridization. First, the slides were dipped into distilled water to remove OCT before a 15-minute bath in 10% neutral buffered formalin (STL286001, Fisher Scientific) (4°C). Next, the slides were bathed for 5 minutes in 50% ethanol (RT), 5 minutes in 70% ethanol (RT), and then 10 minutes in 100% ethanol (RT). The slides were then dried for 5 minutes (RT), and a hydrophobic barrier was applied around each section using an ImmEdge hydrophobic barrier pen. Protease IV was applied to each section for 5 minutes (RT) in the humidity tray (Advanced Cell Diagnostics [ACD], EZ-Batch Slide Holder, Cat # 321716) before beginning the RNAscope Fluorescent v2 assay (ACD, Cat #: 323110) and hybridization to marker probes. RNAscope probes used in this study were as follows: Rn-Nrp1-C1 (ACD, Cat #:1235601-C1), Rn-Calca-C2 (ACD, Cat#:317511-C2), Rn-Nefh-C2 (ACD, Cat#:474241-C2), Rn-Th-C2 (ACD, Cat#:314651-C2), and Rn-Rbfox3-C3 (ACD, Cat#:436351-C3). Probes were incubated for 2 hours, followed by incubation with AMP1 (30 min), AMP2 (30 min), and AMP3 (15 min) before developing the fluorophores. Fluorophores were developed in sequence by incubating in HRP1, HRP2, or HRP3, followed by incubation with TSA vivid fluorophores, and finally followed by incubation with HRP blocker. All incubation steps were completed in the HybEZ oven at 40°C with wash buffer baths in between. The C1 channel (Nrp-1) was labeled with TSA Vivid Fluorophore 570 (ACD, Cat#: 323272), the C2 channel (Calca, Nefh, or Th) was labeled with TSA Vivid Fluorophore 520 (ACD, Cat#: 323271), and the C3 channel (Rbfox3) was labeled with TSA Vivid Fluorophore 650 (ACD, Cat#: 323273). At the end of the RNAscope Fluorescent v2 assay, the slides were coverslipped with VECTASHIELD Antifade Mounting Medium with DAPI (Vector Laboratories, Cat#: H-1200-10), and the edges were sealed with nail polish. Stitched images encompassing an entire DRG section were captured on a Leica DMI8 microscope (Wetzlar, Germany) using a 340 objective and analyzed using QuPath software v0.4.3. Cells with at least 15 puncta associated with an Rbfox3 labeled neuron were considered positive. Each individual dot represents the mean of 3 to 5 quantified sections across 3 DRG per individual rat. The final images were produced in Adobe Illustrator 2022.

### Immunohistochemistry

Mice were euthanized with overdose isoflurane and perfused transcardially with 4% PFA. The L4DRGs and left hind paw skin tissue were dissected and post-fixed in 4% PFA overnight. The DRGs and skin tissues were cryoprotected in 25% sucrose buffer at 4°C, embedded in OCT, sectioned (DRG:14 um and skin: 20um) and mounted on Superfrost Plus slides. Slides were rinsed three times with 0.1M phosphate buffer, blocked in 4% normal donkey serum, 0.25% triton and 1X Phosphate Buffered buffer for 1 hour, and then incubated in primary antibody overnight. For primary antibody incubation, the DRG sections were stained with rabbit anti-mouse CGRP antibody (1:1000, Cat: #198-.2ML, Sigma-Aldrich), and chicken anti-mouse PGP9.5 antibody (1:500, Cat: ab72910, Abcam), while skin sections were stained with rabbit anti-mouse CGRP antibody (1:200, Cat: #198-.2ML, Sigma-Aldrich). Slides were then washed and incubated in secondary antibody for 1 hour. Finally, slides were washed and mounted with prolong golden antifade reagents. Distinct CGRP or PGP9.5 positive neurons were identified using machine learning method in ilastik and further quantified in Fiji software. For quantification of intraepidermal nerve fiber (IENF) density, boundary between epidermis and dermis were identified based on DAPI staining, and CGRP expressing nerve fibers were traced using ilastik software (Figure 5A, right). Then the line grid with area per point 1000 pixels^2 was plotted on the image. The formulation to calculate IENF is as follows: Number of Grids containing epidermis boundary and nerve fiber / Number of grids containing epidermis boundary. At least 3 sections per mice were used to estimate the IENF. The final images were produced in Adobe Illustrator 2025.

### Hind paw injection procedures

Firstly, mice were gently restrained under a fabric cloth and the plantar surface of the left hind paw was exposed. Twenty µL of PBS vehicle or VEGF-A_165_ (4ng, Invitrogen, cat: # RP-8672) was injected intraplantar using an insulin syringe.

### Behavior testing of hindpaw mechanical sensitivity

A Simplified up-down method (SUDO) was performed as described previously to measure mechanical nociception in mouse using von Frey filament. Briefly, mice were placed in acrylic chamber suspended above a wire mesh grid and allowed to acclimate for 1 hour before experiment. The paw withdrawal threshold (PWT) was measured using calibrated von Frey monofilaments von Frey filaments 3 (2.44=0.04g) through 10 (4.31=2g). Von Frey filaments were pressed against the plantar surface of right hind paw until the filament buckled and held for approximately 2 seconds. Flinching upon removal of filament would be considered as a positive response. The test begun with filament 6 (3.61=0.4g) and progressed following an up-down rule that a positive response indicated next lower value filament would be applied in next test, while the negative response indicated a higher value filament applied in the subsequent test. The PWT value was estimated by adding fifth used filament value with an adjustment value of ± 0.5 stimulus intervals (average interval spacing). The average interval spacing according to manufactural value was calculated from slope of value in filament range. The adjustment was positive if the fifth test generated a negative response, or negative if there was a flinching response upon stimulus of fifth von Frey filament. Finally, the PWT value was further normalized by following equation: 10^^PWT value/^10000.

### Behavior testing of abdominal mechanical sensitivity

Mechanical sensitivity of the upper abdomen was assessed using calibrated von Frey monofilaments (North Coast Medical; 0.07 g, 0.4 g). Mice were individually placed in plexiglass chambers mounted on an elevated wire mesh table and acclimated for a minimum of 60 minutes before testing. Each filament was applied to the upper peripancreatic abdominal region, for 1–2 seconds per trial, with 10 trials per filament separated by 30 second inter-trial intervals and 5 minutes between filaments. The percent response was calculated as the number of positive withdrawal responses (hunching, flinching, licking, or guarding) divided by the total number of trials multiplied by 100.

### Cold plantar assay

Mice were allowed to be acclimated in transparent plastic enclosure for at least 1 hour. The top of 3ml BD syringe was removed using razor blade, and then powdered dry ice was pushed into a flattened dense pellet using syringe plunger when open end of syringe was held against glass plate. The dry ice pellet was gently applied to the glass surface underneath the paw of mouse, and the duration of withdrawal latency was recorded. A cutoff of 20s was used to prevent putting paws at risk of cold damage.

### Hindpaw heat sensitivity

The Hargreaves test was designed to measure the thermal sensitivity in mice. Briefly, the mice were placed into transparent chamber on glass plate (temperature held constant at 29°C-30°C) and allowed to acclimate for 1 hour before testing. The heat stimulus generated by light source (intensity: 15%) was applied to the plantar surface of hind paw, and time to withdrawal was recorded. If there is no response by 30 seconds, the test would be terminated, and the thermal withdrawal latency time was recorded as 30 seconds.

### Rotarod

The Rotarod was designed to assess motor function and coordination in rodents. The mice were placed on the rod. Rotarod accelerates 0.5 RPM every 5 seconds, beginning at 2 RPM and maxing out at 60 RPM. The time that mice stayed on the rotating rod before falling was recorded.

### Preparation of dissociated mouse dorsal root ganglion neurons

CGRPcreER:NRP1^fl/fl^:Ai9 mice or CGRPcreER:Ai9 (8 weeks old) from both sexes were euthanized according to institutionally approved procedures. Mice received either corn oil or corn oil containing tamoxifen as desribed above. Dorsal root ganglia from the lumbar region were dissociated as described previously. Briefly, lumbar DRG were collected, trimmed at their roots, and enzymatically digested in DMEM (Cat# 11965, Thermo Fisher Scientific, Waltham, MA) media with neutral protease (3.125 mg/mL, Cat# LS02104, Worthington, Lakewood, NJ) and collagenase type I (5 mg/mL, Cat# LS004194, Worthington, Lakewood, NJ) for 4 min at 37 ⁰C under gentle agitation. The dissociated DRG neurons were gently centrifuged to collect cells and resuspended in complete DRG media (DMEM containing 1% penicillin/ streptomycin sulfate from 10,000 µg/mL stock, and 10% fetal bovine serum (Hyclone)). Cells were seeded on poly-D-lysine-coated 12 mm coverslips and used up to 16 hours after culture.

### Electrophysiological recordings from mouse DRG neurons

Recordings were performed from dissociated DRG neurons obtained from CGRPcreER:NRP1^fl/fl^:Ai9 mice injected with either corn oil (control) or tamoxifen, 14 days previous to euthanasia. Cells were incubated for 30 min with either 0.1% PBS or VEGFA (1 nM) and both were added in the recording solution.

Whole-cell patch clamp experiments were carried out on small-sized DRG neurons from control animals and small-sized DRG neurons from tamoxifen-injected animals, expressing the flourescent protein, tdTomato, using a HEKA EPC-10 patch-clamp amplifier (HEKA Elektronik, Lambrecht, Germany). Borosilicate glass capillaries with resistances between 2.5 and 3.5 MΩ were filled with internal solution containing 120 mM K-Gluconate, 10 mM NaCl, 4 mM Mg-ATP, 5 mM EGTA, 10 mM HEPES, 2 mM MgCl_2_ (Osmolarity= 290 mOsm, pH= 7.3 adjusted with KOH). Action potentials (AP) were evoked in response to depolarizing current injections of 0-120 pA with an increment of 10 pA in 300 ms, in the presence or absence of VEGFA. The bath recording solution contained 130 mM NaCl, 3 mM KCl, 2.5 mM CaCl_2_, 0.6 mM MgCl_2_, 10 mM glucose and 10 mM HEPES (Osmolarity= 320 mOsm, pH= 7.4).

### Data analysis

All graphing and statistical analyses were performed in GraphPad Prism (Version 10). All group data were presented as mean ± SEM. Comparison of two groups were made using unpaired t test. Normality and statistical outlier test were used to confirm appropriate application of parametric statistical tests. Comparisons of more than two groups were made using two-way or three-way ANOVA with repeated measures where appropriate. Pairwise comparisons were made using Tukey’s multiple comparisons test. The details of group size and statistical analysis were further described in the corresponding table and figure legends. No sex differences were detected, so data were collapsed across sexes. Experimenters were blinded to genotype and drug during conduct and analysis of all behavioral assays. Experimenters were blinded to genotype for anatomical analyses.

### Patient and public involvement in research

The study design and progress reports during execution of these studies were presented via poster and oral presentation to the Pancreatic Cancer Action Network at their annual community science summit. Audience participants included multiple stakeholders interested in development of novel pain therapeutics including patients, caregivers, healthcare providers, advocates, and scientists.

## Results

### Neuropilin1 is widely distributed in sensory neurons of mouse and rat

To validate expression of *Nrp1* in sensory neurons, a custom Basescope probe targeted to exon2-3 of *Nrp1* (specifically, the target is the region floxed in the knockout mouse) was combined with immunofluorescence antibody staining for different classical neuron markers in lumbar (L4-L5) dorsal root ganglia (DRG) from adult mice (**Figure 1**). In rat lumbar DRG, RNAscope probes were used to assess co-expression of *Nrp1* across the same subtypes of DRG neurons (**Figure 2**). We observed that 47.67 ± 4.94% of neurons in mouse expressed *Nrp1* mRNA (**Figure 1A, B**). Of the *Nrp1*-expressing neurons, 74.67 ± 5.76% of murine co-expressed neurofilament 200 (NF200) (**Figure 1C**). While tyrosine hydroxylase (Th) is commonly used as a marker of dopaminergic and sympathetic neurons, 18.34 ± 3.65% of DRG sensory neurons expressed Th (**Figure 1A, B**), the majority of which also co-expressed *Nrp1* 76.48 ± 7.02% (**Figure 1C**). While 69.83 ± 5.21% of *Nrp1* positive neurons expressed CGRP, 57.18 ± 10.12% CGRP-expressing DRG neurons also contained *Nrp1* mRNA (**Figure 1C)**. In the rat, a larger proportion of DRG neurons exhibited expression of *Nrp1*, 74.37 ± 1.56% (**Figure 2A, B**). Of those, 86.73 ± 0.75% co-expressed *Nefh (*neurofilament) mRNA. Unexpectedly one-third (34.49 ± 3.2%) of rat DRG neurons also expressed *Th* mRNA, and the majority (69.71 ± 6.56%) of *Th*-expressing neurons co-expressed NRP1 (**Figure 2B, C**). Approximately two-thirds (65.26 ± 7.39%) of *Nrp1* expressing neurons exhibited *Calca* expression. Inversely, 86.77 ± 4.22% of *Calca* positive neurons co-expressed *Nrp1.* These results indicate that while there may be species specific and RNA versus protein variation, NRP1 is widely expressed on both murine and rat DRG sensory neurons, with robust expression in peptidergic neurons.

**Figure 1.**
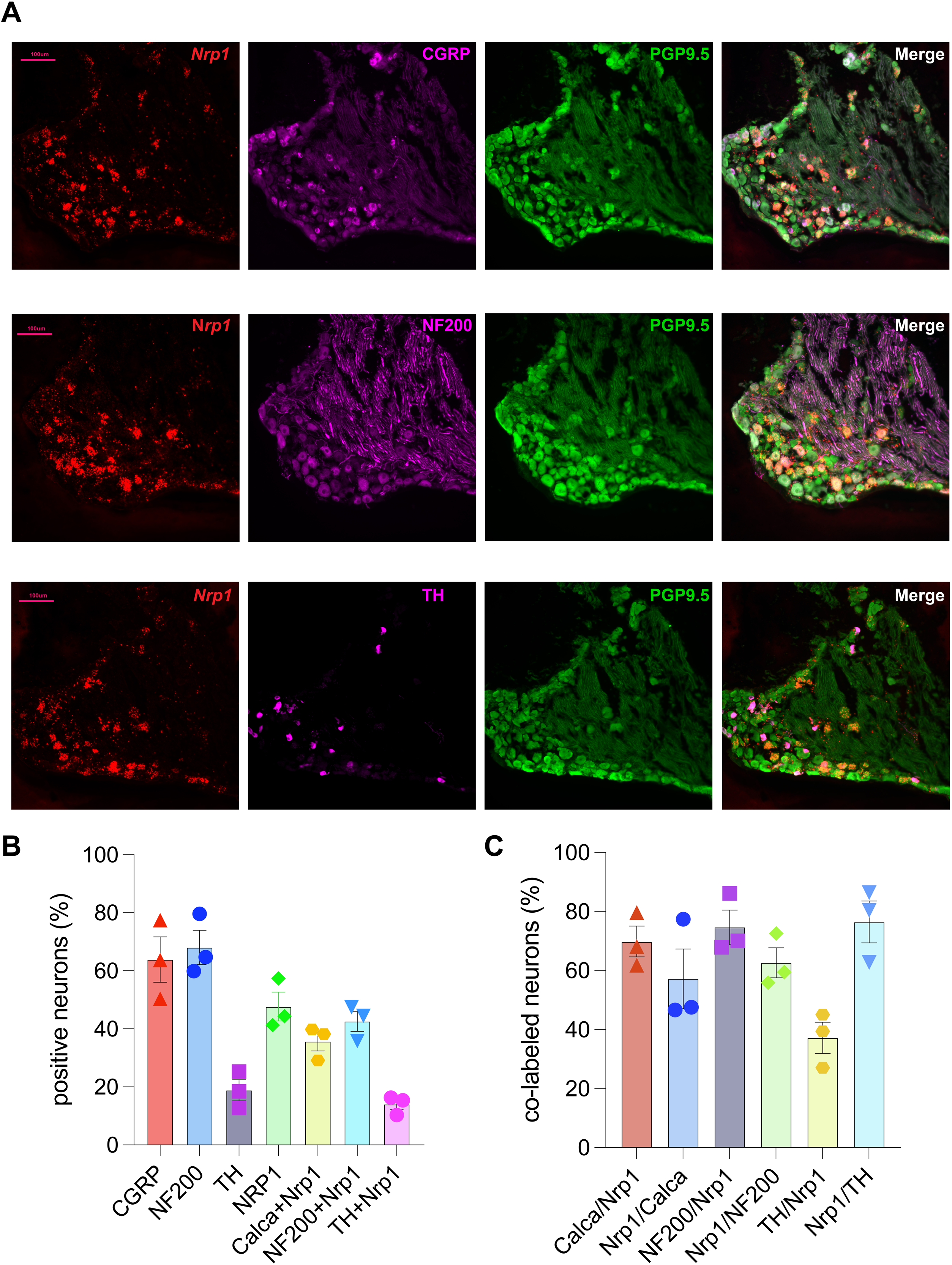
Neuropilin1 is widely distributed in murine spinal sensory neurons. (A) Representative images from fourth lumbar dorsal root ganglion of naïve adult mice illustrating Basescope staining *Nrp1* mRNA combined with immunofluorescent staining for, PGP9.5, CGRP, TH or NF200 protein. Scale bars: 100μm. (B) Bar graph with scatter plots showing the percentages of neuron expressed with distinct markers. (C) Bar graph with scatter plots showing the colocalization between different markers. N=3 mice. Error bars indicate mean ± SEM. Nrp1, neuropilin 1; PGP9.5, protein gene product 9.5; CGRP, calcitonin gene related peptide; TH, tyrosine hydroxylase; NF200, neurofilament 200

**Figure 2.**
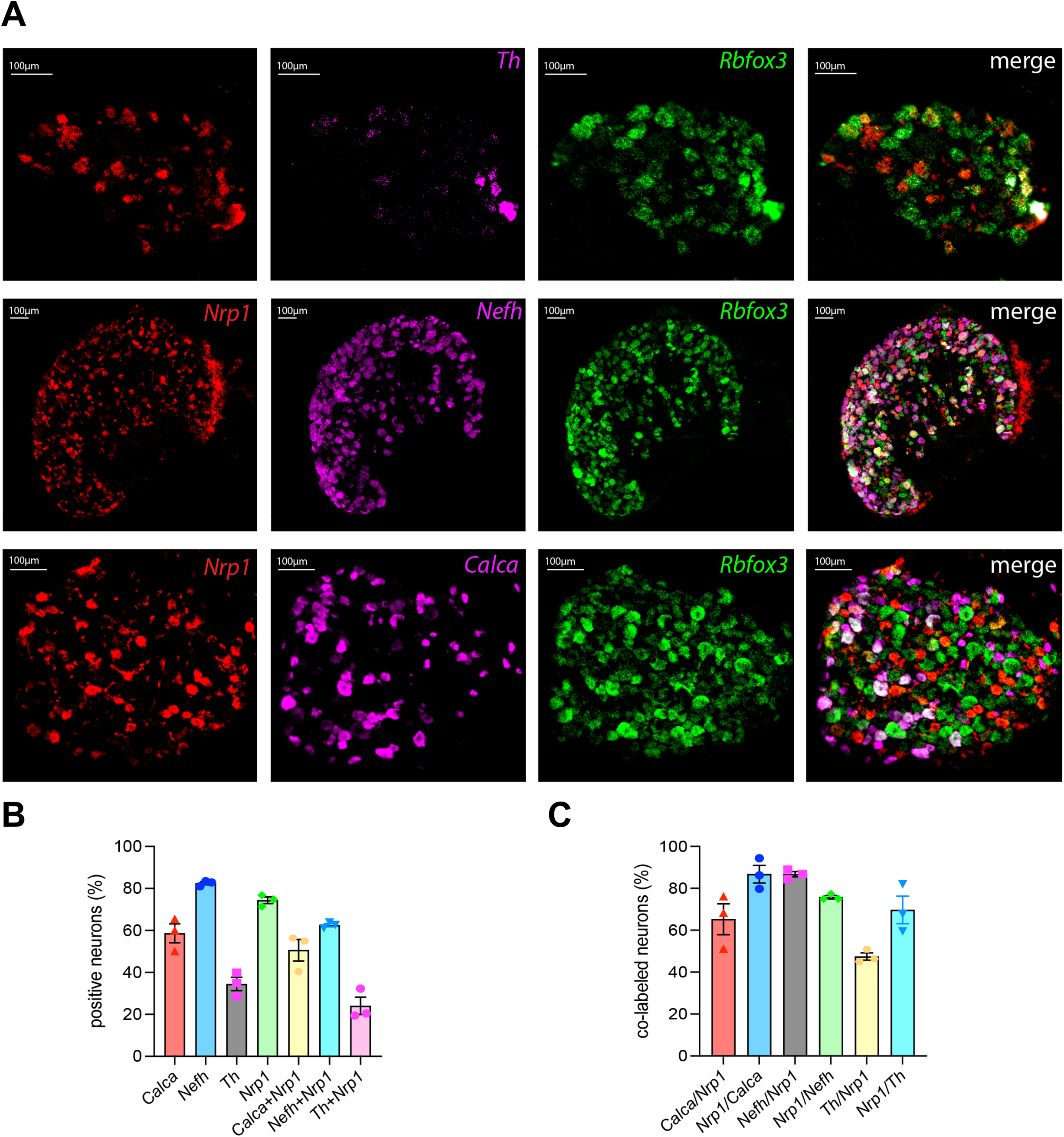
Neuropilin1 is widely distributed in rat spinal sensory neurons. (A) Representative images from lumbar dorsal root ganglion of naïve adult rat illustrating RNAscope staining for *Nrp1, Rbfox3, Calca or Th or Nefh*. Scale bars: 100μm. (B) Bar graph with scatter plots showing the percentages of neuron expressed with distinct markers. (C) Bar graph with scatter plots showing the colocalization between different markers. N=3 rats. Error bars indicate mean ± SEM. *Nrp1,* encodes Nrp1*; Rbfox3,* encodes NeuN; *Calca,* encodes CGRP; *Th* encodes TH; *Nefh*, encodes NF200

### Adult deletion of NRP1 does not alter peptidergic innervation

NRP1 is important in neurodevelopment, particularly axon guidance.[4; 5] Therefore, we utilized CGRPcreER mice to drive NRP1 deletion in adulthood, preserving expression during development. Immunofluorescent staining of L4 DRG was done to confirm that deletion of NRP1 did not alter the total number of CGRP-expressing neurons (**Figure 3**). Compared to genotype control mice (NRP1^fl/fl^), the subset of CGRP-expressing neuron co-expressing NRP1 was significantly decreased in CGRPcreER: NRP1^fl/fl^ mouse (NRP1^fl/fl^ 55.16 ± 3.04% vs CGRPcreER: NRP1^fl/fl^ 14.91 ± 6.76%, P=0.0056), indicating a successful knock down of NRP1 in CGRP-positive peptidergic neurons (**Figure 4**). Importantly, quantification of intraepidermal nerve fiber density confirmed that deletion of NRP1 in adult mouse does not alter CGRP positive innervation of the hindpaw skin (**Figure 5**). The density of CGRP-expressing intraepidermal nerve fibers (IENF) was not different between genotypes, suggesting that any observed phenotype is not due to loss of innervation. Because tamoxifen can cross the blood-brain barrier[38], our manipulation may not have been restricted to the periphery; a limitation of this study is the lack of evaluation of NRP1 expression in the central nervous system.

**Figure 3.**
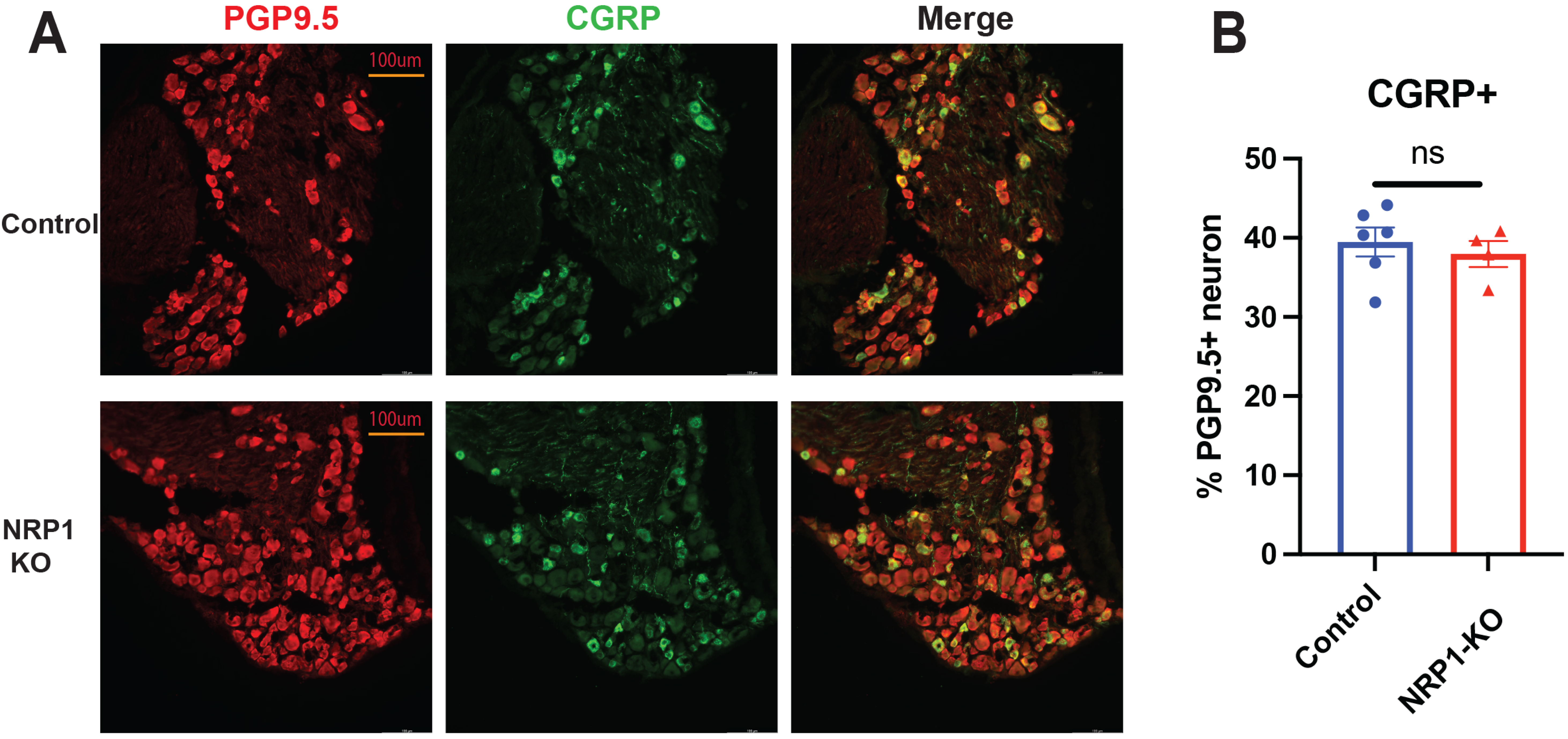
Deletion of NRP1 does not impact CGRP+ neuron survival. (A) Representative images of double label fluorescence for CGRP and PGP9.5 in L4 DRG of NRP1^fl/fl^ and CGRPcreER: NRP1^fl/fl^ mice. Scale bars: 100μm (B) Bar graph with scatter plots shows the percentages of neurons that express CGRP in distinct genotypes. The data are shown in mean ± SEM and analyzed by unpaired t-test. (N=4-5 mice/group, ns: no significance)

**Figure 4.**
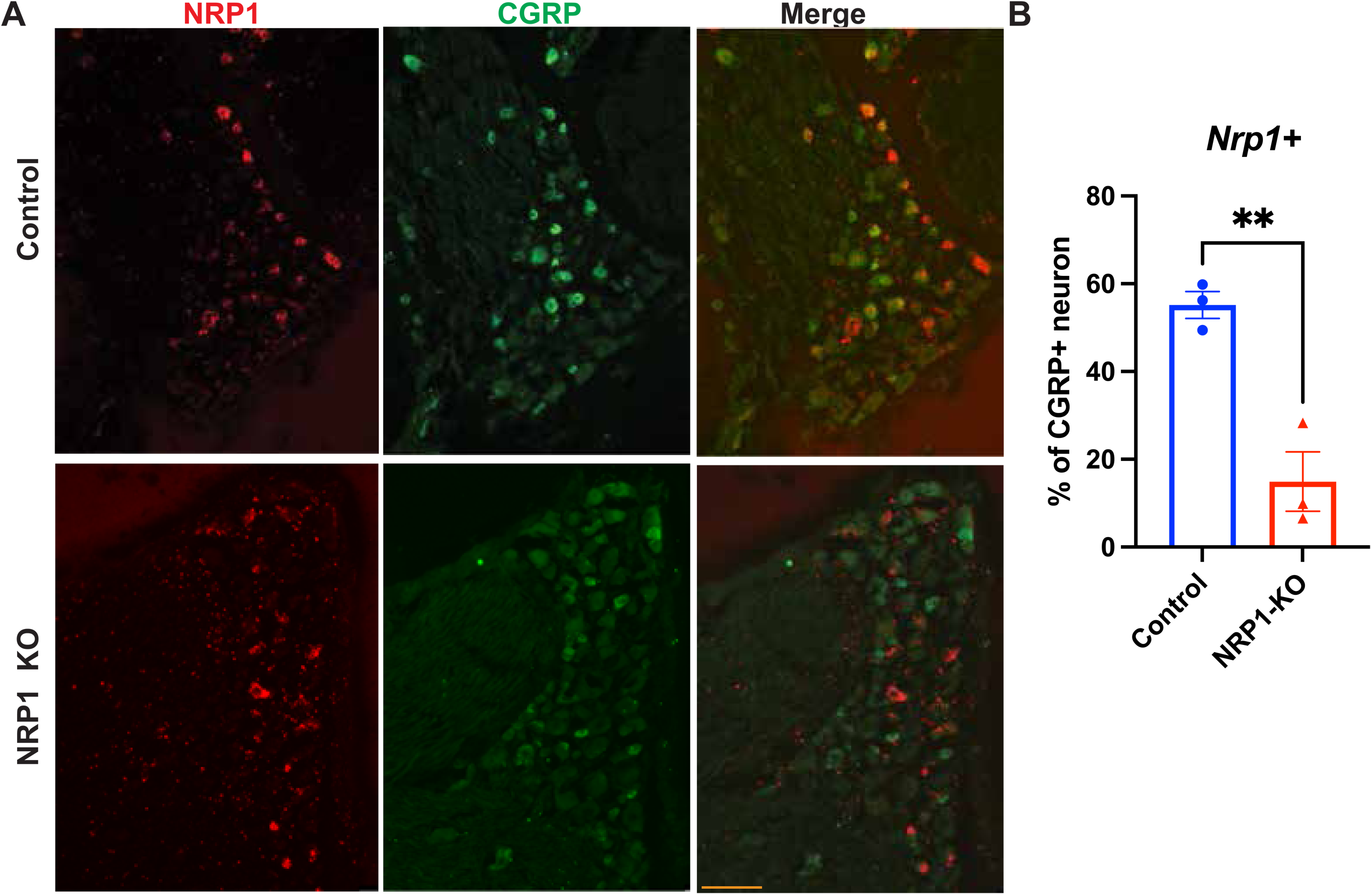
Tamoxifen induces reduction in NRP1 expression in the CGRP+ population of DRG neurons. (A) Representative images of *Nrp1* (Basescope) and CGRP (IHC) expression in L4 DRG from NRP1^fl/fl^ or CGRPcreER: NRP1^fl/fl^ mice after tamoxifen administration. Scale bars: 100μm. (B) Bar graph with scatter plot showing that tamoxifen treatment significantly reduces the number of CGRP+ neurons that express NRP1. N=3 mice. Data were analyzed by unpaired t test, **p < .01. IHC, immunohistochemistry

**Figure 5.**
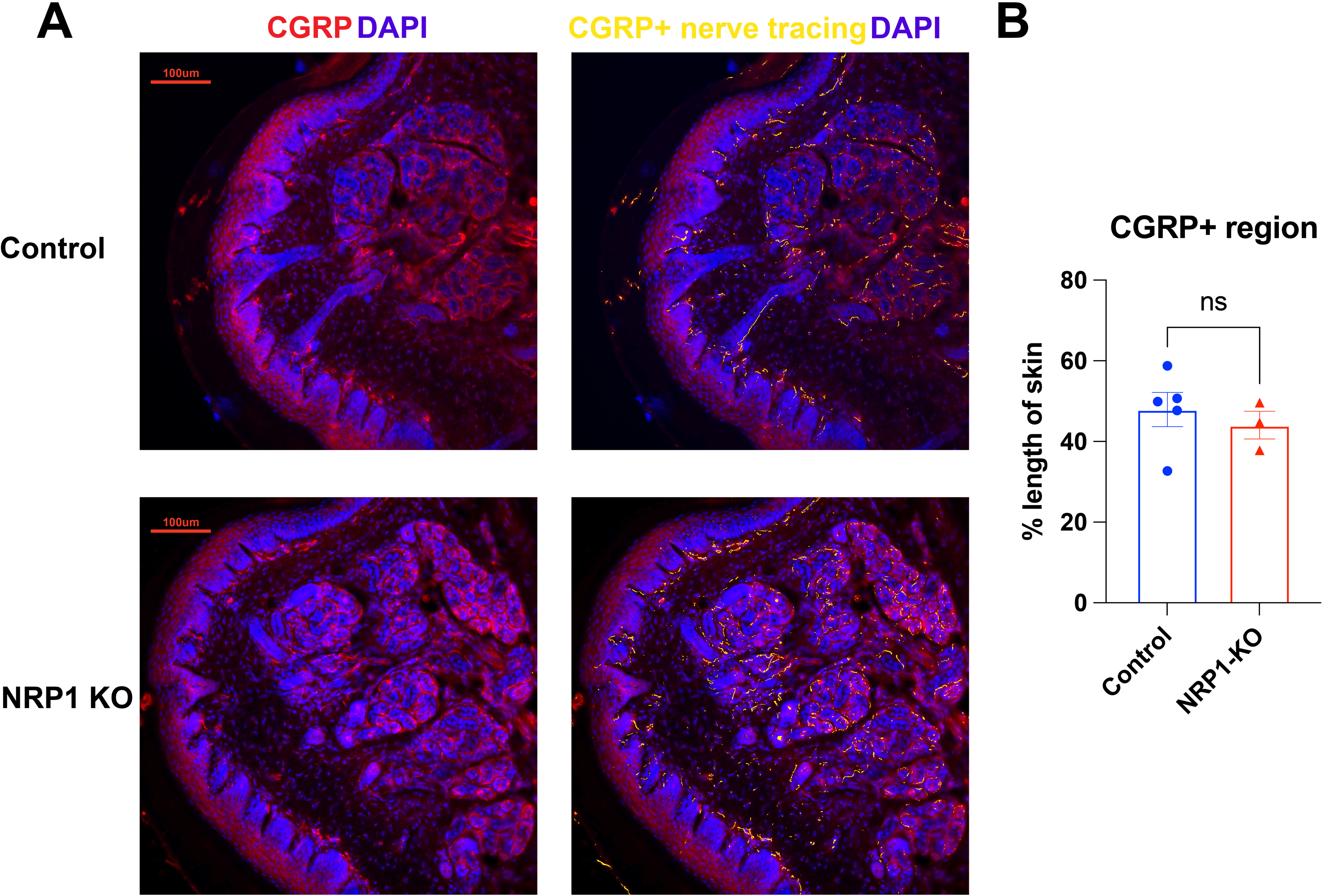
Deletion of NRP1 does not impact CGRP+ cutaneous innervation of the hindpaw. (A) Representative images of immunofluorescence for CGRP in left hind paw papillae of NRP1^fl/fl^ and CGRPcreER: NRP1^fl/fl^ mice. Skin was also labeled with DAPI to identify the border between epidermis and dermis; the CGRP+ nerves (yellow) were traced (yellow) and quantified using ilastik software (shown in right image). Scale bars: 100μm (B) Bar graph with scatter plot shows intraepidermal nerve fiber (IENF) density in distinct genotypes. The data are shown in mean ± SEM and analyzed by unpaired t test. (N=3-5 mice/group, ns: no significance).

### Peptidergic NRP1 mediates mechanical and heat, but not cold, sensitivity

We used standard nociceptive assays to assess the impact of deleting NRP1 from peptidergic sensory neurons. CGRPcreER:NRP1^fl/fl^ mice and littermate controls with or without tamoxifen treatment were assessed for heat sensitivity using the Hargreaves apparatus, hindpaw mechanical sensitivity, and motor function. Mice with loss of NRP1 had significantly higher paw withdrawal latency (PWL) compared to all control groups (**Figure 6A**). Similarly, loss of peptidergic NRP1 resulted in significantly higher mechanical thresholds as measured by the von Frey assay (**Figure 6B)**. Mice with loss of NRP1 had significantly higher paw withdrawal threshold (PWT) compared to all control groups. No sex differences were observed for either heat or mechanical sensitivity (sex*genotype*tamoxifen: hindpaw vf p=0.8283, hargeaves p=0.3890) Cold sensitivity, tested via cold plantar assay, was not impacted by loss of peptidergic NRP1 (**Figure 6C**) as the PWLs to dry ice were not different between any groups. Previous studies have suggested a possible role for CGRP at the neuromuscular junction. Therefore, we also assessed motor function and coordination using the rotarod (**Figure 6D**). There were no significant differences in duration of time spent on the rotarod between genotypes. Given that NRP1 is deleted from all CGRP expressing neurons, the observed hyposensitivity should not be restricted to the hindpaw. Therefore, we assessed abdominal mechanical sensitivity in tamoxifen treated CGRPcreER:NRP1^fl/fl^ mice and littermate controls. At both innocuous (0.07g) and threshold (0.4g) filaments CGRPcreER:NRP1^fl/fl mice^ responded significantly less often compared to controls (**Figure 6E,F**); Like the hindpaw, no sex differences were detected for abdominal mechanical sesntivity (sex*genotype:0.0.07, p=0.6487 and 0.4, p=0.7752).

**Figure 6.**
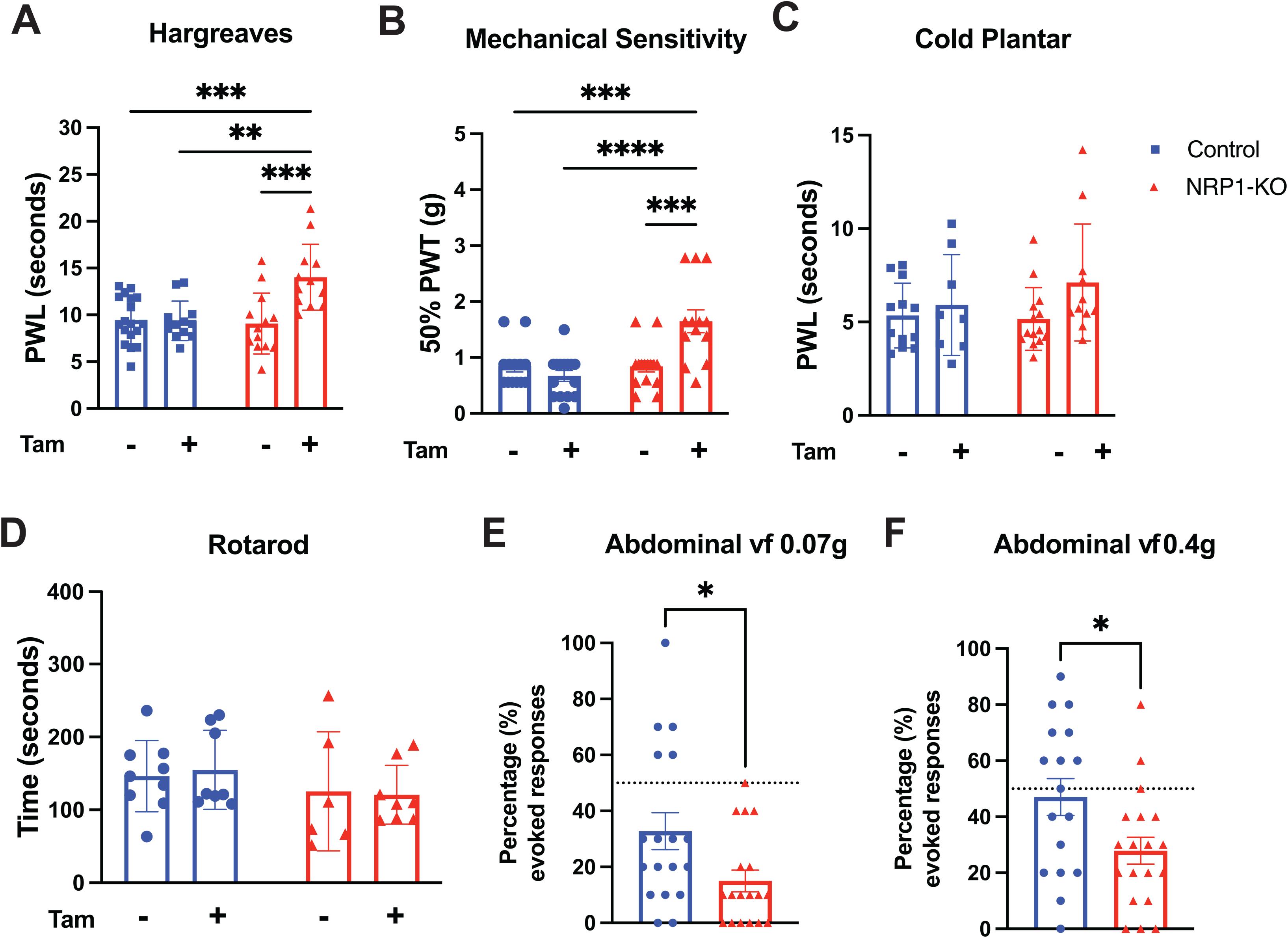
Peptidergic NRP1 regulates heat and mechanical, but not cold, nociceptive behaviors. Behavioral assays were performed on CGRPcreER: NRP1^fl/fl^ or littermate controls with or without tamoxifen treatment. (A) Paw withdrawal latency (PWL) to radiant heat was significantly higher in mice with loss of NRP1 from peptidergic neurons. (B) Loss of peptidergic NRP1 significantly increases mechanical thresholds as measured by von Frey assay. (C) PWL to cold stimulus was not affected by loss of peptidergic NRP1. (D) Locomotion on the rotarod test was not affected by the deletion of NRP1 from CGRP+ neurons. (E) Deletion of NRP1 results in significantly lower response rate to stimulation of the abdomen with an innocuous von frey filament (0.07g) (F) Deletion of NRP1 significantly reduces response rate to 0.4g stimulation of the abdomen. (A-D) Data were analyzed by two-way ANOVA followed by Tukey’s multiple comparisons test. (E-F) Data were analyzed by unpaired t-test. All data are shown in mean ± SEM. *p < .05; **p < .01; ***p < .001; ****p < .0001

### NRP1 is required for VEGFA-dependent hypersensitivity and neuronal sensitization

Binding of VEGFA to VEGFR2/NRP1 receptor complex increases voltage-gated sodium and calcium channel activity and promotes a pain-like phenotype in sensory neuron of rats. To confirm that VEGFA-dependent nociception was disrupted in our mice, we assessed VEGFA-mediated mechanical allodynia. Two weeks after tamoxifen injection in NRP1^fl/fl^ and CGRPcreER: NRP1^fl/fl^ mice, mechanical sensitivity was assayed before, 60 min and 120 min after intraplantar injection of VEGFA or vehicle. Consistent with previous studies, VEGFA induced mechanical hypersensitivity in control mice that was maintained for at least 2 hours (**Figure 7**). However, VEGFA-induced mechanical allodynia was completely prevented by genetic deletion of NRP1 in peptidergic neurons.

**Figure 7.**
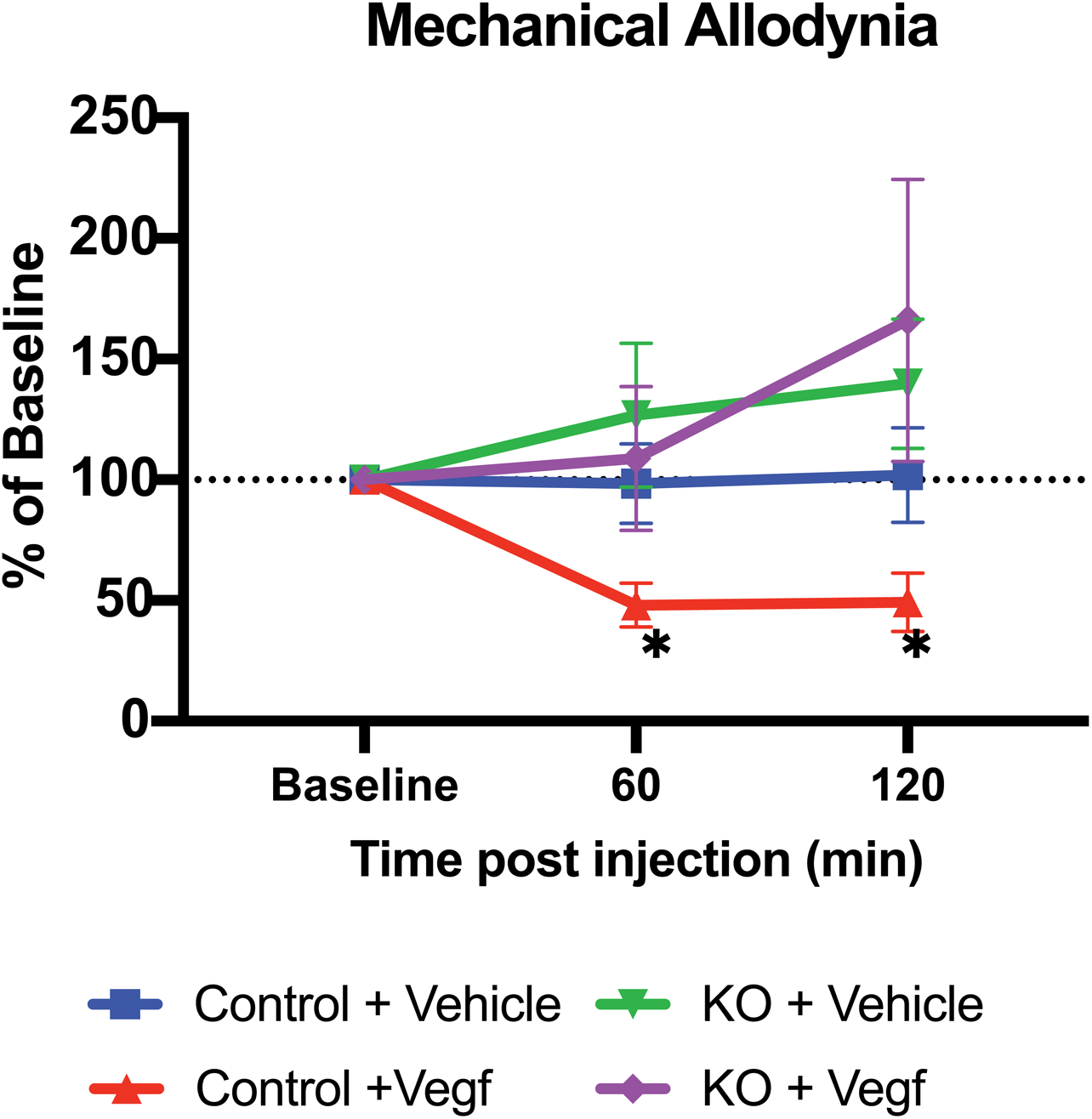
VEGFA induced mechanical hypersensitivity requires peptidergic NRP1. In littermate control mice, intraplantar VEGF-A_165_ (4ng/20μl) induces significant allodynia of the ipsilateral hindpaw. However, VEGF-A_165_ had no effect in peptidergic NRP1 knockout mice, with mechanical thresholds similar to both baseline and vehicle-treatment. N=12-14 mice/group. All data are shown in mean ± SEM. Data were analyzed by three-way ANOVA with repeated measures followed by Tukey’s multiple comparisons test. *p < .05

To assess the role of NRP1 in VEGFA-dependent neuronal sensitization, we performed patch clamp recordings comparing the effects of VEGFA treatment on control and NRP1 knockout neurons. The number of action potentials evoked by current injection alone was not significantly different between control and NRP1 knockout neurons (**Figure 8A, B)**. Rheobase was also not significantly different between neuron genotypes (**Figure 8C**). As anticipated, VEGFA markedly increased the number of action potentials evoked by current injections of 70 pA or greater in control neurons; however, this excitatory effect was completely absent in NRP1-deficient neurons (**Figure 8A, B)**.

**Figure 8.**
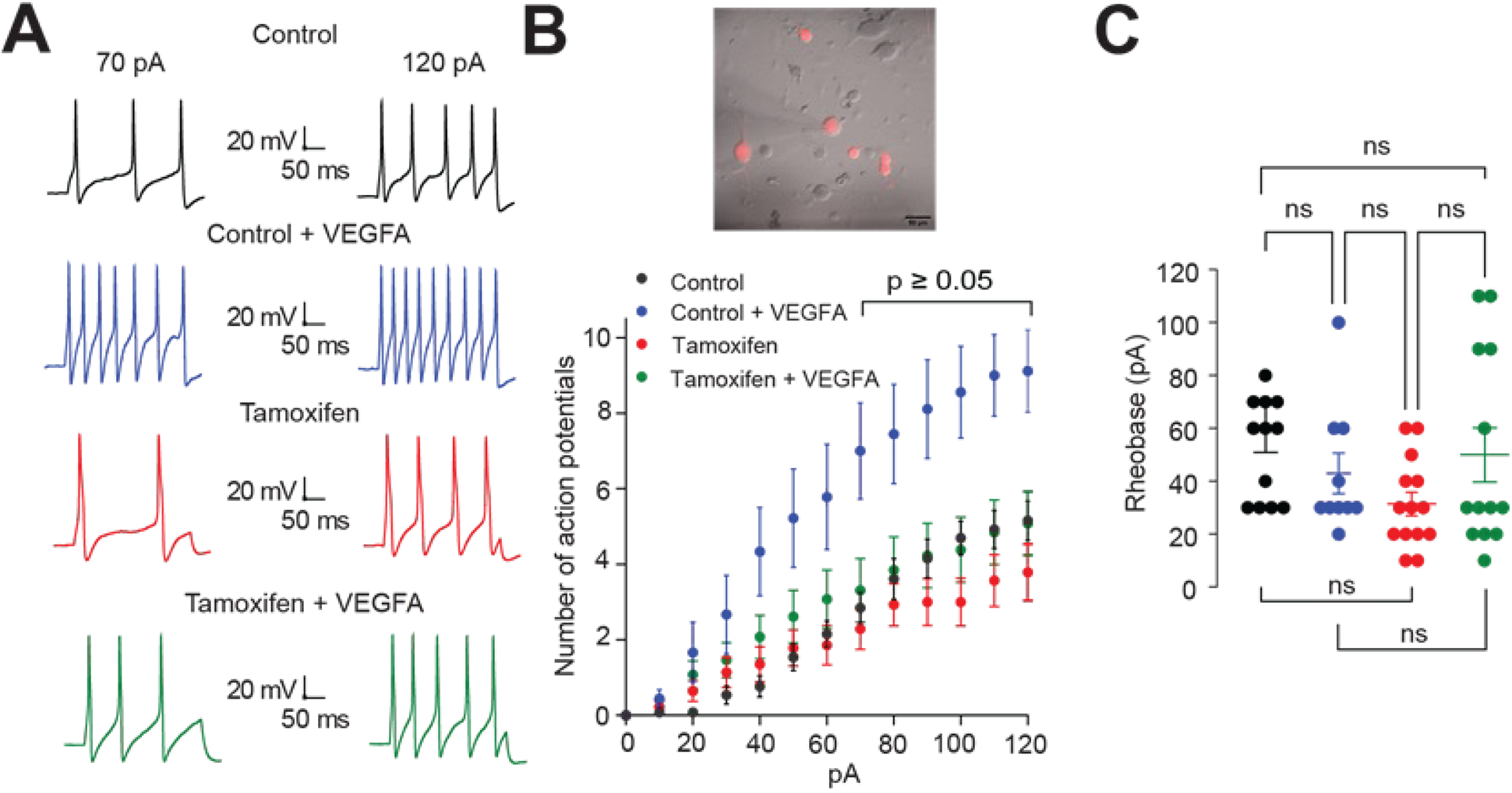
NRP1 is required for VEGFA-dependent nociceptor sensitization. (A). Representative traces of evoked action potentials at 70 pA and 120 pA from dorsal root ganglion neurons (DRGs) from mouse injected with corn oil (black circles), corn oil + VEGFA in the external recording solution (blue circles), tamoxifen (red circles) or tamoxifen + VEGFA in the external recording solution (green circles). (B) Summary of the number of current-evoked action potentials in response to increasing stimuli from 0 to 120 pA of current injection of the indicated conditions. sex: micrography of DRGs from mice previously injected with tamoxifen, red cells were positive for NRP1 conditional KO. **C.** Quantification of rheobase of mouse DRGs treated as indicated. A two-way ANOVA followed by a Tukey’s multiple comparisons test was performed for the 0-120 pA-step excitability protocol, n = 9 to 14 cells per condition. One-way ANOVA followed by a Tukey’s multiple comparisons test was performed for the rheobase data, n = 9 to 14 cells per condition. Data are presented as mean ± SEM.

## Discussion

Previous studies have identified NRP1 as an indispensable co-receptor for VEGFA-mediated pronociceptive actions. Engagement of VEGFA with NRP1/VEGFA receptor complex potentiates the ion channel activities and neurotransmitter release in spinal cord which culminates in occurrence of mechanical allodynia and thermal hyperalgesia. Although NRP1 is widely distributed across DRG neurons, we find that its expression specifically within the peptidergic subset directly regulates basal heat and mechanical nociception. We further demonstrate that the deletion of NRP1 in peptidergic neurons prevents VEGFA-induced hypersensitivity using a transgenic mouse model, concluding that the NRP1 expressed in peptidergic neuron is requisite for the development of VEGFA-induced pain behaviors. Additionally, naïve transgenic mice with deletion of NRP1 in peptidergic neurons had reduced mechanical and thermal sensitivity, indicating that NRP1 is also involved in basal nociception.

Despite a behavioral phenotype in naïve NRP1 knockout mice, we did not detect a robust change in baseline excitability. The hyposensitive behavioral phenotype is consistent with low levels of circulating ligands that, at homeostasis, engage NRP1 signaling pathways and regulate basal nociceptive responses. Indeed, previous studies report low levels of NGF in skin of naïve mice[10; 19]; overexpression of cutaneous NGF promotes hypertrophy of CGRP-expressing cutaneous afferents.[1] Similarly, VEGFA is constitutively released at low levels to provide trophic support keeping tissues, including skin and peripheral nerves, in homeostatic equilibrium.[17; 32; 40] While one might predict that hypoalgesia is a result of hypoexcitability, we did not detect a difference in the two endpoints we measured, ramp-evoked action potentials (AP) and rheobase. The direct effect of deleting NRP1 is to impair functional consequences of ligand binding because, ligands bind NRP1:TRK complexes not ion channels. Without NRP1, membrane TRKs may be nonfunctional. However, ligand binding (and its functional consequences) is also likely to be reduced because of an absence of target receptors. Alternatively, loss of NRP1 may reduce the membrane expression of TrkA, as NRP1 has been reported to function as a chaperone facilitating TrkA trafficking to the cell surface.[25] In either scenario—reduced surface receptor density or a loss of functional intracellular signaling downstream of the NRP1:TrkA complex—the activity of voltage-gated ion channels could be reshaped. Consistent with this, we previously demonstrated that VEGFA, signaling through NRP1, potentiates not only the current density but also the membrane expression of both CaV2.2 (N-type calcium) and NaV1.7 (sodium) channels in primary sensory neurons; pharmacological disruption of the VEGFA–NRP1 interaction with the small-molecule inhibitor NRP1-4 abolished both effects while reducing neuronal excitability and raising the firing threshold. [33] Because CaV2.2 drives calcium-dependent neurotransmitter release, these findings raise the possibility that ligand binding to NRP1:TrkA complexes regulates transmitter release (e.g., CGRP) through enhanced calcium influx, independent of—or in addition to—the triggering of action potentials.

While we focused on CGRP expressing neurons, we confirmed that NRP1 is widely expressed across multiple neuronal subtypes. Previously, Sadighparvar et al., used PIRTcre to delete NRP1 from all sensory neurons and no differences in thermal or mechanical sensitivity were observed in naïve mice.[29] TRPV1cre is a slightly more selective driver, targeting a large heterogeneous population including all C fibers and a subset of A-delta fibers.[6] Similar to PIRTcre:NRP^fl/fl^, we observed no differences in basal somatosensation in the TRPV1cre:NRP1^fl/fl^ mice (Figure S1). There are likely developmental compensatory mechanisms allowing animals to adapt at homeostasis. It is also plausible that the broad loss of NRP1 across multiple neuronal subtypes, some exerting opposing influences, could obscure effects on basal sensitivity. In contrast, selective deletion within peptidergic neurons unmasks a specific role for NRP1 signaling in maintaining sensory homeostasis.

Mammalian nociceptive neurons can be classically categorized as two molecularly and anatomically distinct subsets: peptidergic and nonpeptidergic. Peptidergic neurons express neuropeptides containing Substance P and CGRP and rely on NGF signaling for survival, while nonpeptidergic neurons express IB4 and P2X3, and depend on GDNF signaling for survival. Further, their pain transmission pathways are anatomically segregated in the spinal cord, with peptidergic neurons projecting to lamina I and II_outer_ while non-peptidergic neurons projecting to lamina II_inner._ Our data indicate that NRP1 signaling within peptidergic nociceptors is a key mediator of VEGFA-driven sensitization, as selective loss of NRP1 in these neurons nearly abolishes VEGFA-induced mechanical allodynia and prevents sensitization of CGRP-expressing neurons.

Peptidergic and non-peptidergic neurons may signify the beginning of separate but parallel pain circuits that arise at the level of the spinal cord.[3] Non-peptidergic neurons terminating on lamina II interneurons target limbic brain areas via lamina V projection neurons, contributing to the affective aspects of pain, whereas the pathway for peptidergic sensory neurons is largely through lamina I projection neurons, which contributes to the sensory-discriminative aspects of pain.[3] Although electrophysiological studies indicate that most nociceptors are polymodal neurons that respond to multiple noxious stimulus modalities, distinct subsets of nociceptors selectively mediate behavioral responses to different noxious stimuli applied to the skin. Mrgprd+ neurons, which constitute most of the cutaneous nonpeptidergic neurons, selectively respond to noxious mechanical stimuli but not to noxious heat.[7] In contrast, CGRP expressing sensory neurons are necessary for transmission of noxious heat but unnecessary for response to mechanical stimuli. Further, IB4-negative neurons have much larger heat activated inward currents compared to IB4-positive neurons. Together, the above suggests that peptidergic neurons are more important in transduction of noxious heat stimuli while non-peptidergic neurons are responsible for relaying noxious mechanical information. In our behavior tests on naïve mice, loss of NRP1 in peptidergic neurons impacts both thermal and mechanical sensitivity: CGRPcreER:NRP1^fl/fl^ mice displayed significantly increased thermal PWLs and mechanical PWTs compared to other control groups, suggesting that NRP1 expressed in peptidergic neurons is necessary to maintain normal thermal and mechanical sensitivity. While previous studies suggest that peptidergic neurons are dispensable for cutaneous mechanical pain transmission, CGRP is expressed in both c and Aδ mechanosensitive neurons. Unfortunately, our von frey assay for mechanical sensitivity may not be adequate to determine the impact on this population which would be better assessed by the pinprick assays. Regardless, our von frey results indicate a novel role of NRP1 in peptidergic neurons in processing noxious mechanical stimuli. It is likely that subsets of peptidergic neurons encompass multiple distinct signaling pathways, and NRP1 expressing peptidergic neurons define a subset of peptidergic signaling that promotes transmission of noxious mechanical pain.

In conclusion, our study demonstrates that loss of NRP1 expression in peptidergic neurons reduces noxious heat and mechanical sensitivity as well as prevents VEGFA-dependent neuronal sensitization and hypersensitivity. NRP1 is overexpressed in sensory neurons following nerve injury[13]. In addition to the VEGF family, NRP1 is a co-receptor for multiple TRK ligands, many of which are also known to drive pro-inflammatory and pro-nociceptive associated with neuronal sensitization and pain, including NGF[8; 30; 43; 44], EGF[27], IGF[11; 39], PDGF[2] and TGFß[16; 42]. Given that loss of NRP1 in our hands was sufficient to promote multimodal hyposensitivity, it may be a useful therapeutic target for painful conditions in which one or more pro-nociceptive mediators are being released near NRP1-expressing nerve terminals. Thus, targeting NRP1 expressed in peptidergic neurons may provide a precise and reliable target for pain management.

## Acknowledgements

NCI R01 CA285585 (JLS), Pancreatic Cancer Action Network 22-20-SALO (JLS), NINDS RF1 NS131165 (RK), NINDS T32 NS073548 (OLB). Central South University, Xiangya School of Medicine provided partial stipend and bench-fee for SX (PIs: Margaret McDonald, Xiaochuang Wu). No federal U.S. funds were paid to foreign components. The authors have no additional disclosures or conflicts of interest. All data are available upon reasonable request to the senior authors.

---

**Figure S1.**
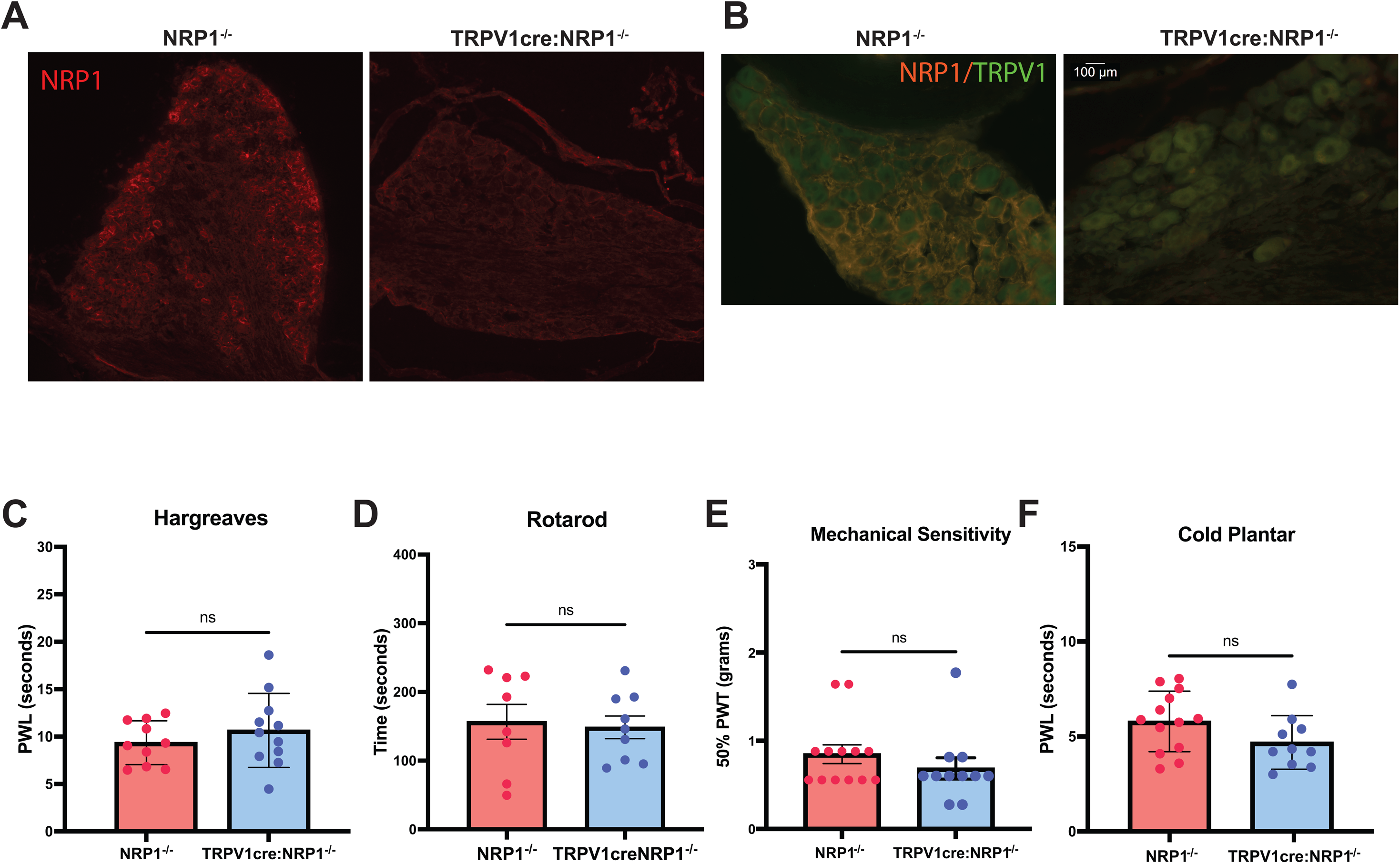
TRPV1creNRP1fl/fl mice did not display attenuated mechanical and thermal sensitivity. 10X (A) and 20X (B) example micrographs of NRP1 in L4 DRG from control (NRP1^−/−^) and TRPV1cre:NRP1^−/−^mice. rHargreaves (C), Rotarod assay (D) hind paw Von Frey assay (E) and cold plantar assay (F) were performed to compare motor function and sensitivity to various noxious stimulus between NRP1fl/fl and TRPV1creNRP1fl/fl mice. Paw withdrawal threshold, Paw withdrawal latencies and duration of time on accelerating rod Data were measured and shown as mean ± SEM; Data were analyzed by unpaired t test. (ns represent no significance)

**Supplementary Table 1.**

| **Antibody** | **Host** | **Concentration** | **Catalog number** | **RRID** |
| --- | --- | --- | --- | --- |
| Anti-mouse CGRP | Rabbit | 1:200(skin);<br>1:1000 (DRG);<br>1:500 (Basescope) | Sigma C8198 | AB_259091 |
| Anti-mouse PGP9.5 | Chicken | 1:500 | Abcam 72910 | AB_1269734 |
| Anti-mouse NF200 | Rabbit | 1:400 | Abcam AB8135 | AB_306298 |
| Anti-mouse TH | Rabbit | 1:500 | Sigma AB152 | AB_390204 |
| Anti-Rabbit Cy3 | Donkey | 1:500 | Jackson<br>ImmunoResearch<br>711-165-152 | AB_2307443 |
| Anti-Chicken AF488 | Donkey | 1:500 | Jackson<br>ImmunoResearch<br>703-545-155 | AB_2340375 |
| Anti-Rabbit Cy5 | Donkey | 1:500 | Jackson<br>ImmunoResearch<br>711-175-152 | AB_2340607 |

